# Inflammatory cytokine profile and plasticity of brain and spinal microglia in response to ATP and glutamate

**DOI:** 10.1101/2020.11.02.365676

**Authors:** Sam Joshva Baskar Jesudasan, Somnath J Gupta, Matthew A Churchward, Kathryn Todd, Ian R Winship

## Abstract

Microglia are the primary cells in the central nervous system that identify and respond to injury or damage. Such a perturbation in the nervous system induces the release of molecules including ATP and glutamate that act as damage-associated molecular patterns (DAMPs). DAMPs are detected by microglia, which then regulate the inflammatory response in a manner sensitive to their surrounding environment. The available data indicates that ATP and glutamate can induce the release of pro inflammatory factors TNF (tumor necrosis factor), IL-1β (interleukin 1 beta) and NO (nitric oxide) from microglia. However, non-physiological concentrations of ATP and glutamate were often used to derive these insights. Here, we have compared the response of spinal cord microglia (SM) relative to brain microglia (BM) using physiologically relevant concentrations of glutamate and ATP that mimic injured conditions in the central nervous system. The data show that ATP and glutamate are not significant modulators of the release of cytokines from either BM or SM. Consistent with previous studies, spinal microglia exhibited a general trend towards reduced release of inflammatory cytokines relative to brain-derived microglia. Moreover, we demonstrate that the responses of microglia to these DAMPs can be altered by modifying the biochemical milieu in their surrounding environment. Preconditioning brain derived microglia with media from spinal cord derived mixed glial cultures shifted their release of IL-ß, IL-6 and IL-10 to a less inflammatory phenotype consistent with a spinal microglia.

## Introduction

Microglia have the capacity to respond to pathogens, insults, and injuries that disrupt the homeostasis of the CNS (1). They are able to respond to a wide variety of environmental stimuli due to their repertoire of receptors that can sense damage-associated molecular patterns (DAMPs) and pathogen-associated molecular patterns (PAMPs) in the central nervous system (CNS) (2, 3). Adenosine triphosphate (ATP) and glutamate are examples of DAMPs that are released into the extra cellular milieu in response to various injuries or perturbations of CNS, including cell death due to stroke, spinal cord injury, traumatic brain injury (2,3). ATP and glutamate are well known for their role as chemotactic agents that recruit microglia to the site of injury (4, 5). ATP has also been shown to induce upregulation of pro inflammatory factors such as TNF (tumor necrosis factor), IL-1β (interleukin 1 beta) and NO (nitric oxide) by microglia through purinergic receptor pathways (1,6). Similarly, glutamate has been shown to induce pro-inflammatory factors such as TNF, IL-1β, and NO in microglia through glutamate receptors (1,7,8). However, most studies that investigated these DAMPs used specific receptor agonists or antagonists or used non-physiological concentrations (≥10 fold above the physiological concentrations) to determine their effects on microglia in culture (9,10). ATP and glutamate can activate ionotropic [ATP: P2×1-7, glutamate: AMPA(Glu2/3), kainate (Glu5)] and metabotropic receptors (ATP: P2Y1-6,11-14, glutamate: mGlu1-8) on microglia. Previously, it was shown that 1 mM glutamate induces release of TNF through AMPA (GluR2-4) and kainate (GluR5) receptors (11). The Group II mGlurR2 and 3 specific agonist DCG-IV also induces TNF release by microglia (12). Interestingly, selective inhibition of group II mGlu5 reduces TNF release by LPS-activated microglia (13). In the present study, physiologically relevant concentrations of ATP and glutamate that mimic injury were used to test if they have a differential effect on brain-derived (BM) and spinal cord-derived (SCM) primary microglia. Previous studies in models of rat ischemia and TBI have shown that glutamate concentrations of up to 10 µM occur in the uninjured CNS extracellular milieu, and that this concentration is increased to ≥30 µM after ischemic injury or TBI (9,14,15,16). Extracellular glutamate concentrations of more than 100 µM induce neurotoxicity (15). While there is no general consensus on ATP concentration in the extracellular milieu during homeostasis, it has been suggested that after an injury a 500 µM concentration and above occurs (17,18) and several studies have utilized a 1mM concentration of ATP to measure the effect of ATP on microglia *in vitro* (19,20, 21,22,23). Microglial phenotypes are dependent on region of origin (24) age, sex and environment (25,26,27). Notably, spinal cord microglia (SCM) have a reduced inflammatory profile in response to activation by lipopolysaccharide (LPS) relative to microglia derived from the brain (BM) (28). However, the responses of SCM (relative to BM) to physiological stimulation with ATP and glutamate have not been investigated. Given that previous data suggest a less inflammatory phenotype in SCM, this study tested the hypotheses that physiological activators such as ATP and glutamate would induce a reduced inflammatory profile in SCM compared to BM. Previously, we have shown that microglia from different regions of brain (hippocampus and thalamus) that are exposed to conditioned media from striatum acquired an inflammatory profile similar to that of microglia originally derived from the striatum (23). This suggests that microglia are a highly plastic population of cells. Hence, we also hypothesized that regional heterogeneity is not fixed and that BM exposed to SCM condition media would move toward an inflammatory profile similar to that of SCM.

To test the first hypothesis, BM and SCM from postnatal Sprague-Dawley rat pups were activated in vitro with ATP (1 mM) in vitro as previously established (23) and glutamate (10 µM to model physiological concentration in rat brain parenchyma, 30 µM to mimic the concentration measured in vivo in rat brain parenchyma after ischemic or TBI injury, and 100 µM to mimic excitotoxic injury) (29,9,14,15,16). To test the second hypothesis, BM were incubated in conditioned media from brain and spinal cord mixed glia (BMix CM and SMix CM, respectively) to replicate the extracellular environment in which BM and SCM were cultured. BM conditioned in BMix cm and SMix cm were activated with ATP, glutamate (10 µM, 30 µM, 100 µM) and LPS (1 µg/ml) and pro-inflammatory factors released were measured.

## Results

### Secretion of pro-inflammatory effectors by BM and SCM exposed to ATP

ATP is commonly released in the extracellular milieu after injury to the CNS (1,2,6). To investigate phenotypic differences between SCM and BM in response to this endogenous activator, isolated SCM and BM microglia were treated with ATP. Previous studies suggest that ATP concentrations of 1 mM are sufficient to induced a pro-inflammatory profile (NO-Nitic oxide, TNF-Tumor necrosis factor, IL-1β - Interleukin 1β and IL-6 - Interleukin 6) in BM (19,20, 21,22,23). BM and SCM were therefore treated with 1 mM ATP, and pro-inflammatory factors (NO, TNF, IL-6, IL-1β) released into cell culture media were measured using the Greiss assay (NO) and ELISAs (TNF, IL-6, IL-1β) (Figure 1, Table 1). Two-away ANOVA significant interaction between microglia and treatment for NO, TNF, IL-1β, a significant main effect of treatment for release of NO, TNF, IL-1β and a significant main effect of microglial origin was observed for TNF, IL-6 and IL-1β, the data is summarized in table 1. Sidak post-hoc test revealed that NO, TNF and IL-1β released by LPS (1 µg/ml) activated SCM LPS was significantly less that of BM LPS (BM LPS vs SCM LPS: NO p = 0.042, TNF p < 0.001, IL-1β p<0.0001). No other significant comparisons were found, suggesting that group differences were largely driven by differential responses to LPS activation rather than ATP treatment.

**Table 1:** Two-way ANOVA details for effect of ATP on SCM and BM

| Pro inflammatory Effectors | ATP |  |  |  |  |  |
| --- | --- | --- | --- | --- | --- | --- |
|  | Interaction |  | Treatment |  | Microglia |  |
|  | F (Dfn, Dfd) | p | F (Dfn, Dfd) | p | F (Dfn, Dfd) | p |
| TNF | (2, 42) = 20.005 | p<0.0001 | (2, 42) = 78.349 | p<0.0001 | (1, 42) = 23.196 | p<0.0001 |
| IL-6 | (2, 30) = 0.5 | p=0.6119 | (2, 30) = 17.816 | p<0.0001 | (1, 30) = 7.531 | p=0.0101 |
| IL-1β | (2, 30) = 9.951 | p=0.0005 | (2, 30) = 3.372 | p=0.0478 | (1, 30) = 26.051 | p<0.0001 |
| NO | (2, 24) = 4.119 | p=0.0290 | (2, 24) = 14.577 | p<0.0001 | (1, 24) = 2.900 | p=0.1015 |

**Figure 1.**
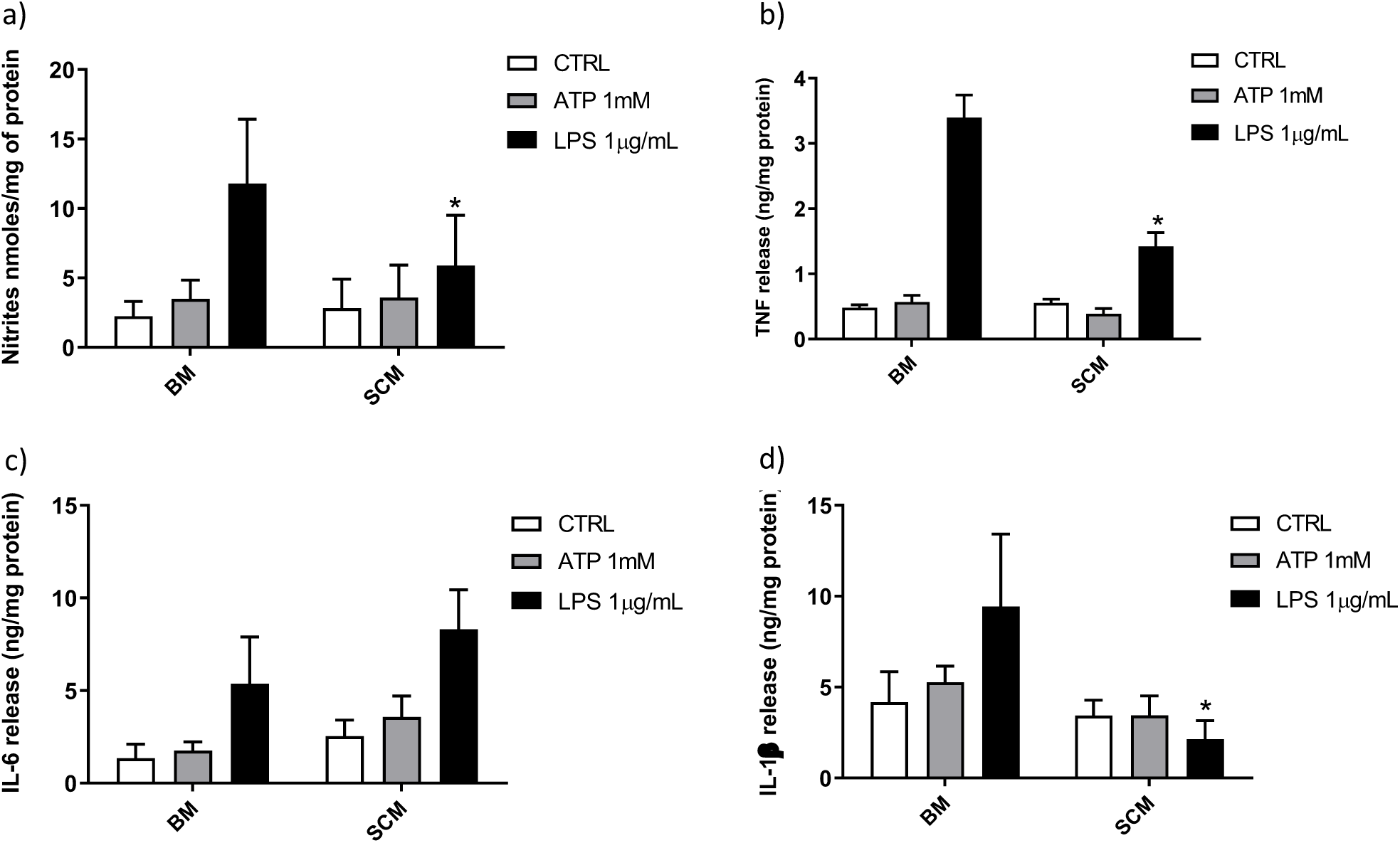
Release of the pro-inflammatory effectors by ATP activated BM and SCM. **(a)** Two-away ANOVA identified significant interaction between microglia and treatment for NO and a main effect of treatment for NO. However, Sidak post-hoc test revealed a significant difference between BM LPS and SCM LPS, * represents p < 0.05, n = 5where n represents the number of independent experiments, where an independent experiment is a separate microglia preparation, **(b & c)** Two-away ANOVA identified significant interaction between microglia and treatment TNF and IL 6. A significant difference between BM LPS and SCM LPS was revealed by the Sidak post-hoc test, * represents p < 0.05, n = 8 for TNF and n = 6 for IL6 where n represents the number of independent experiments, where an independent experiment is separate microglia preparation, **(d)** Two-away ANOVA identified significant interaction between microglia and treatment for IL-1β and a significant main effect of microglia. A Sidak post-hoc test revealed a significant difference between BM LPS and SCM after LPS treatment. * represents p < 0.05 for comparison, n = 6 where n represent the number of independent experiments, where an independent experiment is a separate microglia preparation. ATP treatment did not induce a significant difference in the release of inflammatory factor NO, TNF, IL 6, IL −1 β between BM and SCM ATP treatments. Bars represent mean ± s.e.m.

### Secretion of proinflammatory effectors by BM and SCM exposed to glutamate

Glutamate agonists have been shown to induce pro- or anti-inflammatory profile in a receptor dependent manner in microglia (9,10). However, direct study of microglia phenotype in response to physiological concentrations of glutamate has not been frequently investigated. Here, 10 µM (representing physiological levels of glutamate in CNS extra cellular milieu), 30 µM (levels present in *in vivo* ischemic injury), 100 µM (levels present at excitotoxicity injury sites) (29,16) were selected as concentrations for treatment of BM and SCM (Figure 2). Two-way ANOVA identified significant interaction between microglia and treatment for NO, IL-6, IL-1β and TNF. A significant main effect of treatment for TNF, IL-6, and IL-1β and significant main effects of microglia for IL-1β, IL-6 and NO were also detected (Table 2). Sidak post-hoc analysis revealed that NO released by SCM LPS was significantly less than that of LPS treated BM (BM LPS vs SCM LPS p < 0.0001, Figure 2a), and that LPS mediated IL-6 and IL-1β release by SCM was significantly less than that of BM (BM LPS vs SCM LPS p < 0.05) (Figure 2c and Figure 2d). While a qualitative trend towards reduced release of all inflammatory mediators in response to glutamate in SCM was suggested, post-hoc comparisons did not identify statistically significant differences in cytokine release between BM and SCM in response to different glutamate concentrations.

**Table 2:** Two-way ANOVA details for effect of glutamate on SCM and BM

| Pro inflammatory Effectors | Glutamate |  |  |  |  |  |
| --- | --- | --- | --- | --- | --- | --- |
|  | Interaction |  | Treatment |  | Microglia |  |
|  | F (Dfn, Dfd) | p | F (Dfn, Dfd) | p | F (Dfn, Dfd) | p |
| TNF | (4, 30) = 1.302 | p=0.2916 | (4, 30) = 38.127 | p<0.0001 | (1, 30) = 1.954 | p=0.1725 |
| IL-6 | (4, 40) = 2.075 | p=0.1022 | (4, 40) = 16.869 | p<0.0001 | (1, 40) = 3.484 | p<0.0001 |
| IL-1β | (1, 40) = 1.426 | p=0.0005 | (4, 40) = 20.53 | p<0.0001 | (1, 40) = 7.232 | p=0.0104 |
| NO | (4,20) = 13.13 | P<0.0001 | (4,20) = 50.43 | P<0.0001 | (1,20) = 15.67 | p=0.0008 |

**Figure 2.**
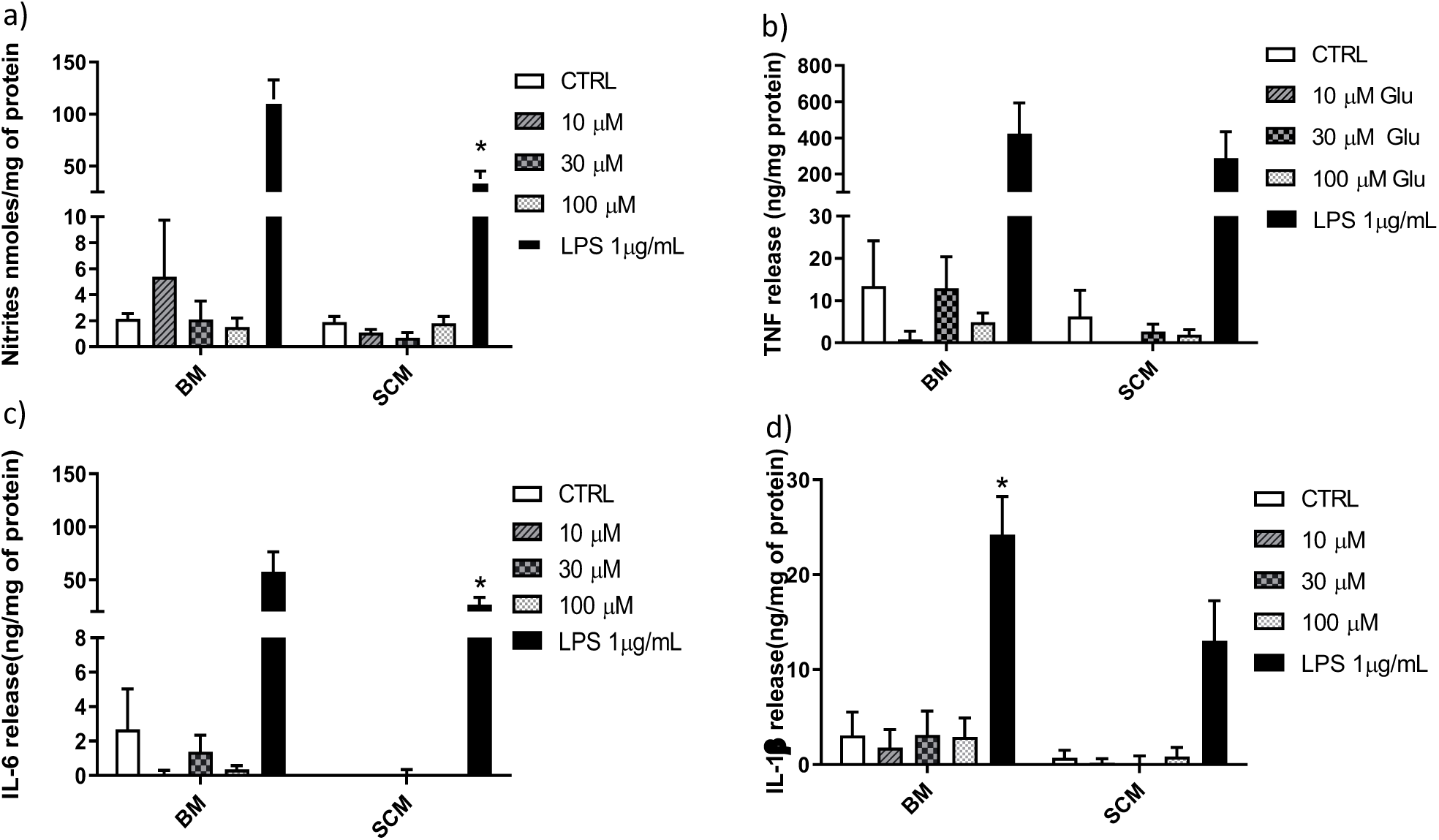
Release of the pro-inflammatory effector by BM and SCM exposed to glutamate. **(a)** Two-way ANOVA identified significant interaction between microglia and treatment for NO. There was also a significant main effect of microglia and treatment, n = 3 where n represent the number of independent experiments, where an independent experiment is a separate microglia preparation. **(b & c)** Two-way ANOVA revealed a significant main effect for glutamate treatment for TNF. n= 4 for TNF and n = for IL6 where n represent the number of independent experiments, where an independent experiment is a separate microglia preparation, **(d)** Two-way ANOVA revealed a significant main effect for microglia and treatment, n= 5 where n represent the number of independent experiments, where an independent experiment is a separate microglia preparation. * represents p < 0.05 a significant difference between respective LPS treatments revealed by the Sidak post-hoc test. Bars represent mean ± s.e.m.

### Secretion of proinflammatory effectors by BM exposed to BMix and SMix CM

In addition to regional heterogeneity in microglia isolated from different areas of the brain it has been shown that the microglia phenotypes differ with age and that phenotypes from different age groups are not fixed. Notably, conditioned media from microglia derived from one age group as well as a different region can be used to alter the phenotype of microglia derived from animals of a different age group or region of brain (23). This suggests that the BM are highly plastic cells capable of adapting to immediate environment and their phenotype is dictated not only by genetics but also their immediate environment. Therefore, the pro-inflammatory profile of SCM may be due to the conditioning by spinal cord mixed glia culture media. We hypothesized that conditioning BM with SMix CM (spinal cord mixed glia conditioned media) would alter the concentrations of inflammatory factors released by BM. To test the hypothesis, BM were incubated in SMix CM or BMix CM (brain mixed glia conditioned media) prior to treatment with ATP, glutamate, or LPS. BM incubated in BMix CM and SMix CM were activated with glutamate (10 µM, 30 µM, 100 µM), ATP (1 mM), LPS (1µg/ml) and the release of IL-6, IL-10 and IL-1β was measured. Two-way ANOVA indicated that conditioned media and treatment had a significant main effect for IL-6 and IL-1β release, and a significant interaction between media and treatment condition for IL-10 (Figure 3, Table 3). Overall, the data suggest that BM release more IL-6 and IL-10, and less IL-1β when incubated with SMix media relative to BMix media. Sidak post-hoc tests suggest that IL-6 released in response to LPS was significantly greater in BM incubated in SMix CM (BMix CM BM LPS vs SMix CM BM LPS, p < 0.0032) (Figure 3a). Moreover, BM release of IL-6 and IL-10 in response to 100 µM glutamate was significantly greater when incubated in SMix CM (BMix CM BM 100 µM vs SMix CM BM 100 µM, p = 0.0480, p = 0.0283 respectively) (Figure 3a & 3c) and suggested an overall reduction in IL-1β release with SMix CM (Figure 3b). Sidak post-hoc comparisons did not reveal any significant difference between individual BMix CM and SMix CM treatment groups for IL-1β (Figure 3b).

**Table 3:** Conditioned media-mediated cytokine profile of BM and SCM

| Pro inflammatory Effectors | Condition Media |  |  |  |  |  |
| --- | --- | --- | --- | --- | --- | --- |
|  | Interaction |  | Treatment |  | Condition Media |  |
|  | F (Dfn, Dfd) | p | F (Dfn, Dfd) | p | F (Dfn, Dfd) | p |
| IL-10 | (5, 42) = 3.026 | p=0.0295 | (5, 24) = 1.544 | p=0.2139 | (1, 24) = 3.262 | p=0.0835 |
| IL-6 | (5, 24) = 2.487 | p=0.0597 | (5, 24) = 92.45 | p<0.0001 | (1, 24) = 12.98 | p=0.0014 |
| IL-1β | (5, 24) = 0.440 | p=0.8161 | (5, 24) = 12.37 | p<0.0001 | (1, 24) = 6.965 | p=0.0144 |

**Figure 3.**
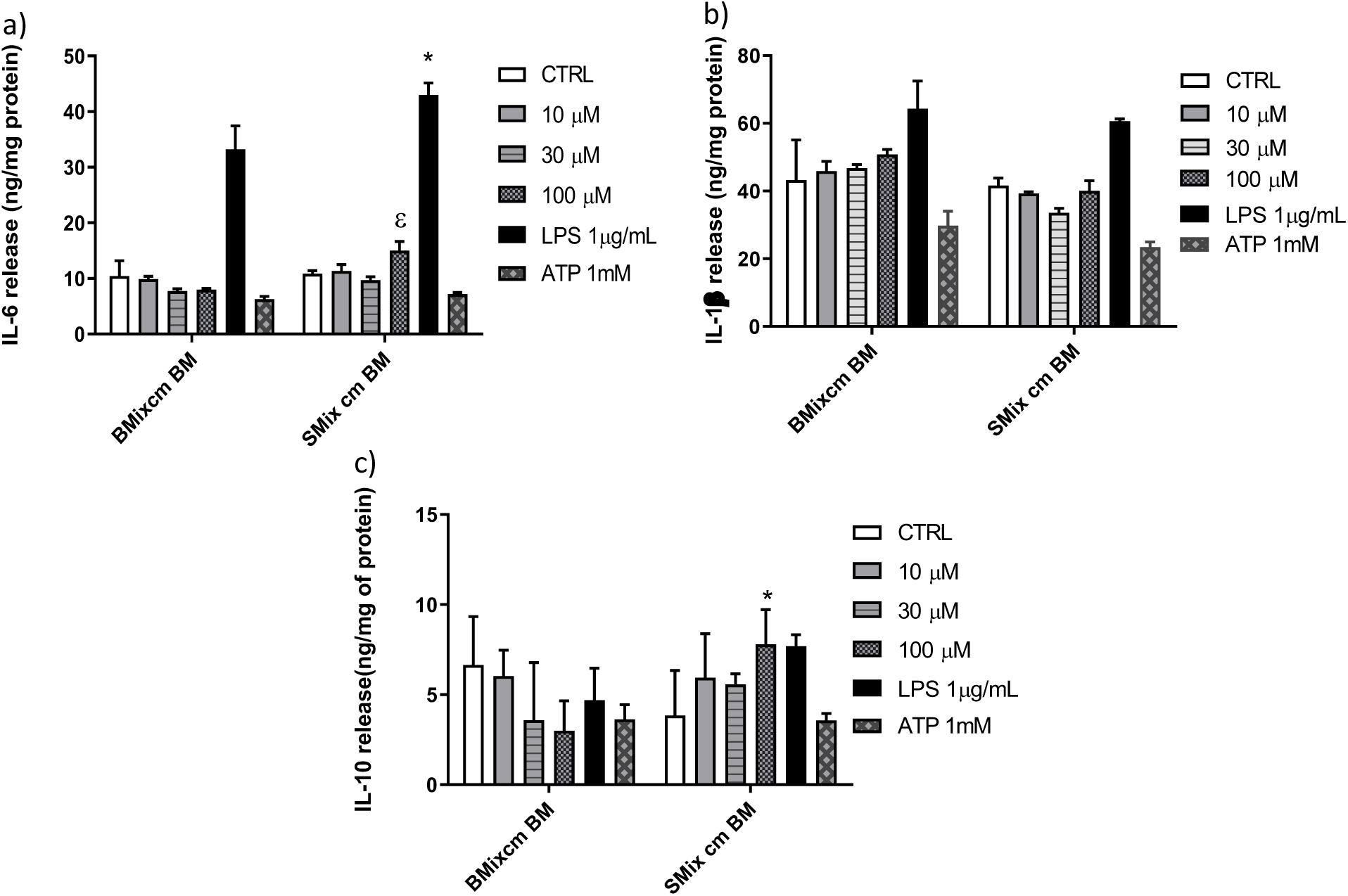
Secretion of the pro-inflammatory cytokines by BM exposed to BMix CM and SMix CM. **(a)** Two-way ANOVA also revealed a significant main effect of conditioned media and treatment for IL-6 release. ε represents significant difference for 100 µM glutamate treatment (p = 0.0480) between brain mixed glia conditioned media (BMix CM) and spinal mixed glia conditioned media (SMix CM) * represents p < 0.05 significant difference between LPS treatment for BMix CM and SMix CM as revealed by the Sidak post-hoc test, n = 3 where n represent the number of independent experiments, where an independent experiment is a separate microglia preparation, **(b)** Two-way ANOVA revealed a significant main effect of conditioned media and treatment for the release of IL 1β. However, the Sidak post-hoc test did not reveal any significant differences between BMix CM and SMix CM. n= 3 where n represents the number of independent experiments, where an independent experiment is a separate microglia preparation. Bars represent mean ± s.e.m., **(c)** Two-way ANOVA revealed a significant main effect of interaction. Sidak post-hoc test showed a significant difference (p = 0.0295) for 100 µM glutamate treatment between brain mixed glia conditioned media (BMix CM) and spinal mixed glia conditioned media (SMix CM). n= 3 where n represents the number of independent experiments, where an independent experiment is a separate microglia preparation. Bars represent mean ± s.e.m.

## Discussion

### ATP and glutamate activation of microglia

Insults to the CNS, such as ischemic stroke and spinal cord injury, cause neuronal injury and lead to release of neurotransmitters such as ATP and glutamate into the extracellular milieu (1, 8, 6, 16). Increased glutamate concentration in the extracellular milieu induces excitotoxicity injury to neurons and chemotaxis of microglia to the site of injury (1,5,8). Similarly, ATP released in the extracellular milieu induces microglial activation as well as chemotaxis of microglia to the site of injury (26, 6). However, the response of microglia to known activators and to perturbations in the CNS are also dependent on their immediate environment (31).

### ATP mediated cytokine profile of BM and SCM

Previous studies suggest that SCM have a reduced inflammatory phenotype in response to LPS exposure relative to BM. In this study, we found no differences in the release of the proinflammatory factors NO, TNF, IL-6 or IL-1β between BM and SCM treated with ATP (1mM) (Figure 1 a-d). This data suggests regional heterogeneity (i.e. microglia from brain vs microglia from spinal cord) does not influence ATP (purinergic receptors) mediated activation of microglia, unlike LPS where the SCM inflammatory profile was reduced compared to that of BM (a result replicated here). This data supports the inference that ATP by itself may have a primary role as a chemoattractant rather than an immunomodulator (33,34).

Moreover, previous studies have also shown that combination of DAMPs (ATP+LPS) induces a much more rapid induction of inflammatory cytokine IL-1β than just LPS (peaked at five hours post treatment vs at 25 hours) (35). The data from this study indicates that origin of microglia (brain or spinal cord) does not alter their inflammatory behaviour in response to ATP alone. ATP may instead act in combination with other DAMPs to evoke induction of inflammatory cytokine release after an injury, in addition to a role as a chemoattractant.

### Glutamate mediated cytokine profile of BM and SCM

Previous studies shown that 1 mM glutamate induced release of TNF through AMPA (GluR2-4) and kainate (GluR5) receptors (12). Similarly, Group II m GlurR2 and GlurR3 specific agonist DCG-IV induce TNF release by microglia (12). However, 1 mM glutamate does not mimic the concentration of glutamate at the site of an ischemic injury or TBI (9,14,15,16). The purpose of this glutamate study was to test if glutamate can induce the release of inflammatory factors at physiological concentrations mimicking ischemic and excitotoxicity injury, and to test if glutamate could induce differential release of inflammatory factors by BM and SCM. Glutamate at physiological concentrations did not induce significant release of pro-inflammatory factors (TNF and IL-6) and there was no significant difference between BM and SCM release in individual treatment groups (Figure 2 b-c). However, an overall reduction in NO and IL-1β release by SCM was observed in glutamate experiments (Figure 2a and Figure 2d). Interestingly, it has been shown that glutamate can induce chemotaxis in microglia (5). Thus, glutamate at physiological concentrations may also play a primary role as a chemoattractant in addition to its role as mediator of inflammation. At higher concentration of glutamate, the BM and SCM did not show any significant change in release of proinflammatory factors (TNF, IL-6 and NO). One possible explanation could be related to the ability of microglia to uptake glutamate and convert it to glutathione, thus protecting from oxidative stress (36). Glutamate treatment did not show any morphological change in microglia, suggesting that glutamate is not directly involved in initiating inflammatory response from microglia (37).

### Conditioned media-mediated cytokine profile of BM and SCM

All microglia in the CNS originate from the yolk sac during embryogenesis. This suggests that all microglia in CNS are genetically very similar (38). However, microglia are very versatile and studies have shown that they can have different phenotypes (surface receptor expression, pro-, and anti-inflammatory factors release) in distinct regions of brain (23). For example, the P2XR7 and P2YR12 expression is higher in striatum than in other regions of the brain (23). This suggests that microglia adapt to their immediate environment, potentially due to paracrine signaling from neighbouring cells that alters the extracellular milieu.

The conditioned media experiments were designed to answer the question whether the BM exposed to SMix CM would adopt a SCM-like phenotype compared to BM exposed to BMix CM. Notably, SMix CM significantly increased release of IL-6 compared to BMix CM (Figure 3a), suggesting that the environmental milieu can significantly affect phenotype for certain inflammatory molecules. This observed increase in release of IL-6 is consistent with the prior evidence of increased LPS induced release of IL-6 from SCM compared to BM (28), and may result from increased levels of GMCSF or IFN γ levels in SMix CM that can potentiate the release of IL-6 (39). Similarly, an overall reduction in IL-1β release was observed after incubation of BM with SMix CM, paralleling observations of SCM treated with LPS (28). An increase was observed for the release of anti-inflammatory cytokine IL-10 in SMix CM after glutamate treatment, which suggests that SMix CM can induce the release of IL-10 compared to BMix CM and favour a switch from a proinflammatory phenotype to anti-inflammatory or neuroprotective phenotype (40, 41). The conditioned media experiments therefore provide further evidence that BM phenotype is plastic and can be modified by varying the extracellular environment, thus supporting the hypothesis that not only the region of origin but also the immediate environment determine the phenotype of microglia. This further supports the postulate that microglial phenotype is not fixed by region of origin and that immediate environment plays a crucial role in the diversity of microglial phenotypes after perturbation or injury to CNS. The data also aligns well with previous studies that demonstrated that the inflammatory profile of microglia is dependent on the severity of injury, where microglia were beneficial to outcome of mild injury (26,31).

This study investigated whether ATP and glutamate induce a differential inflammatory profile in SCM vs. BM and addressed whether these mediators at physiological concentrations induce the release of inflammatory cytokines by BM and SCM. Our study is one of the few studies to examine the effects of physiological concentrations of glutamate on microglia. Our results suggest that ATP and glutamate do not induce significant release of pro-inflammatory factors such as NO, TNF, IL-6 and IL-1β. However, our data do not preclude a role for these activators in the inflammatory response to brain injury. Both ATP and glutamate are involved in chemotaxis of microglia and can potentially recruit microglia to the site of injury or perturbation (20, 5, 34). Moreover, previous findings demonstrated that ATP in combination with LPS induced faster maturation and release of intracellularly accumulated IL-1β (35), suggesting a modulatory role that was not tested here. Earlier studies showed that exposing microglia from thalamus and hippocampus to conditioned media from striatum induces a striatum-like phenotype in the microglia (23). Similarly, we found that SMix CM altered the response profiles of BM compared to BMix CM, including increased release of IL-6 and reduced release of IL-1β in response to LPS activation and increased release of IL-10 and IL-6 following glutamate treatment (Figure 3 a-c). Overall, our data suggest that ATP and glutamate at physiological concentration do not induce a inflammatory cytokine release in BM and SCM and provide further support that the phenotype of BM as well as SCM in vitro is determined by both region of origin and immediate environment in addition to other factors such as age and sex (25, 26, 27).

## Methods

### Media and reagents

Hanks Balanced Saline solutions (HBSS), Dulbecco’s Modified Eagle Medium – Hams’F12 nutrient mixture (DMEM-F12), DMEM-F12 with HEPES (DMEM-F12/HEPES), 0.25% trypsin-EDTA, fetal bovine serum (FBS), and Penicillin-Streptomycin (P/S) were from Gibco (ThermoFisher Scientific, Burlington, ON). ATP, glutamate, lidocaine HCl, Triton X-100, LPS, and sodium nitrite standard solution were from Sigma (Oakville, ON).

### Primary mixed glia preparation

All animal protocols were conducted in accordance with Canadian Council on Animal Care Guidelines and approved by the Animal Care and Use Committee: Health Sciences for the University of Alberta. Brains and spinal cords for establishing primary mixed glial cultures were obtained from postnatal day one or two male Sprague-Dawley (SD) rat pups as previously described (30,31). The SD rat pups were euthanized, their brains (four) and spinal cords (twenty) were dissected and placed in dissection buffer (HBSS with 200 U/mL penicillin, 200 μg/mL streptomycin). Meninges and blood vessels were removed under a dissection microscope. Tissues were cut into small pieces and incubated in 0.25% Trypsin-EDTA for 25 min at 37°C, and collected by centrifugation (2000 x g, 2 min). Trypsin was inactivated with maintenance media (DMEM/F12 supplemented with 10% FBS and 200 U/mL penicillin, 200 μg/mL streptomycin) and tissues were dissociated by trituration in maintenance media and centrifuged at 2000 x g for 2 mins. The brain and spinal cord cell pellets were re-suspended in maintenance media and seeded at equal density into cell culture-treated T75 flasks coated with poly-L-lysine. Cells were maintained in a 37°C, 5% CO_2_ humidified incubator with maintenance media replaced twice weekly.

### Microglia isolation by lidocaine HCl isolation method

Microglia were isolated from primary mixed glial cultures at 21 days in vitro by modification of the lidocaine HCl shaking method (31,32). At 24-hours prior to isolation cultures were refreshed with DMEM/F-12 supplemented with 10% FBS, and immediately before isolation this medium was collected, filtered (0.22 µm), and diluted 1:1 with DMEM/F-12 to make conditioned medium. Brain and spinal mixed glial cultures at 21 days in vitro were refreshed with DMEM-F12/HEPES with 10% FBS at 37°C 30 mins before lidocaine HCl treatment. Lidocaine HCl was added to the primary mixed glial culture to a final concentration of 15 mM in the media, the cultures were incubated for 3 mins and shaken for 7 mins in an orbital shaker at 50 rpm at 37°C. Microglia were collected from the media by centrifugation (2000 x g, 2 minutes at room temperature). Cell pellets were gently re-suspended in 1 mL of conditioned media using 1 mL pipette tip. Cell suspensions were washed by adding 9 mL of their respective conditioned media and centrifuged at 2000 x g for 2 mins at room temperature. The cell pellets were again re-suspended by trituration in their respective conditioned media and the cells were counted and seed at a density of 1×10^5^ cell/mL in 48 or 24 well poly-L-lysine coated polystyrene plates. The microglia were allowed to settle in the plates for 10 mins at 37°C in a 5% CO_2_ incubator after which non adherent cells were washed gently with DMEM at 37°C and respective conditioned media were added to the BM and SCM and the mixture were allowed to recover overnight. Conditioned media were replaced with fresh DMEM/F-12 immediately prior to treatments.

### Nitric Oxide (NO)

NO release was measured indirectly by quantifying the stable metabolite nitrite in culture media using a method described by Griess, (1879). Media were collected 24 hours after treatment, 100 µL of treatment or control media were added to a 96-well plate in duplicates followed by 50 µL each of 1% sulphanilamide (in 3N HCl) and 0.02% N-napthylethylenediamine per well. Absorbance was read at 540 nm and the amount of nitrite metabolite was interpolated from a set of standards measured in parallel.

### Enzyme Linked Immunosorbent assays (ELISA)

Commercial ELISA kits were used to measure TNF, IL-1β, IL-6 and IL-10 in media (DuoSet, R&D Systems Minneapolis, USA). ELISA procedures were carried out according to manufacturer protocols.

### Statistics

Statistical analyses were carried out using two-way ANOVA followed by Sidak’s methods to test for significance between treatment groups. n represents a single independent experiment (i.e. an independent culture preparation) with a minimum of three technical replicates. Each technical replicate represents a well in a 12, 24 or 48 well culture plate. All statistical analyses were done using Graphpad Prism version 8.3.0.

## Data availability

All data are contained within the article.

## Funding and additional information

This work was supported by the Canadian Institutes of Health Research (IRW), the Heart and Stroke Foundation of Canada (IRW), the Natural Sciences and Engineering Research Council (IRW), and the George Davey Endowment for Brain Research (KGT).

